# Single molecule tracking and analysis framework including theory-predicted parameter settings

**DOI:** 10.1101/2020.11.25.398271

**Authors:** Timo Kuhn, Johannes Hettich, J. Christof M. Gebhardt

**Affiliations:** Institute of Biophysics, Ulm University, Albert-Einstein-Allee 11, 89081 Ulm, Germany

## Abstract

Imaging, tracking and analyzing individual biomolecules in living systems is a powerful technology to obtain quantitative kinetic and spatial information such as reaction rates, diffusion coefficients and localization maps. Common tracking tools often operate on single movies and require additional manual steps to analyze whole data sets or to compare different experimental conditions. We report a fast and comprehensive single molecule tracking and analysis framework (TrackIt) to simultaneously process several multi-movie data sets. A user-friendly GUI offers convenient tracking visualization, multiple state-of-the-art analysis procedures, display of results, and data im- and export at different levels to utilize external software tools. We applied our framework to quantify dissociation rates of a transcription factor in the nucleus and found that tracking errors, similar to fluorophore photobleaching, have to be considered for reliable analysis. Accordingly, we developed an algorithm, which accounts for both tracking losses and suggests optimized tracking parameters when evaluating reaction rates. Our versatile and extensible framework facilitates quantitative analysis of single molecule experiments at different experimental conditions.

## Introduction

Single-molecule experiments are gaining increasing importance when investigating dynamical and structural parameters such as binding kinetics, diffusion coefficients or spatial distributions of biomolecules in living systems. In these experiments, the biomolecule of interest is typically fused to a fluorescent label, such that the signal of fluorescent photons in successive recordings reports on the position and movement of the biomolecule. Extracting quantitative information from such movies includes linking individual detections of biomolecules at consecutive time points to continuous tracks ^1–6^. In recent years, several tracking algorithms have been adapted to tracking of biomolecules, including basic nearest neighbour and more complex algorithms such as Kalman filtering, combinatorial optimization, multiple hypothesis tracking or neural networks^2,7–11^. Still, the intuitive application of tracking algorithms is challenged by a significant dependence on empirical parameters, such as the tracking radius which is usually determined on visual aspects ^5,12^. Furthermore only few of the particle tracking and analysis softwares published up to now were made accessible to a broader audience by providing an intuitively operable graphical user interface ^9,13–17^. In addition, the subsequent analysis steps to extract quantitative information from tracked molecules are mostly left to additional software such as SMTracker ^18^, Spot-ON ^19^ or vbSPT ^20^. Thus, analyzing whole data sets consisting of multiple movies or comparing different experimental conditions is cumbersome as they require extensive manual intervention. Overall, there is high demand for a user-friendly, comprehensive program covering tracking, analysis and data visualization of single molecule experiments.

Linking detections of biomolecules into tracks unavoidably comes with errors, amongst others due to missed detections, inappropriate tracking radius or mix-up at high molecule density ^2,5,12^. The probability for errors can be reduced by low molecule densities, though ^21^. Premature loss of a track also occurs if the fluorescent label photobleaches. Photobleaching can be conveniently corrected for, e.g. by comparison with immobile histone molecules ^22^, by ensemble measurements ^23,24^, or by time-lapse imaging ^25^. In contrast, errors inherent to the linking process are more challenging to tackle and often only assessed qualitatively. Recently, the effect of allowing several gaps in assembling tracks of immobile molecules has been considered ^17^. Also the impact of the tracking radius on the diffusion coefficient has been discussed ^5,12^. In diffusion analysis, molecules diffusing out of the focal plane need to be accounted for ^19^. However, a theoretical description of tracking errors and how they can be corrected for is missing for tracking immobile fluorescently labelled biomolecules.

We introduce TrackIt, an integrative tracking and analysis software for fast and extensive analysis of single molecule data sets within a single framework. Our user-friendly graphical user interface (GUI) provides access to two different tracking algorithms and multiple analysis procedures for kinetic and spatial parameters. Moreover, it comes with several data visualization options. Additionally, tracking results can be exported to utilize external analysis algorithms such as Spot-ON and vbSPT ^19,20^. We organize analysis parameters and associated data in a batch structure such that the workflow starting from spot detection over tracking to quantitative analysis can be repeated in a single step. This enables convenient comparison of different tracking parameters as well as of data sets recorded at different experimental conditions. Furthermore, we introduce a formalism to estimate tracking errors when linking immobile molecules and quantify the decrease of tracking losses if a gap frame is allowed. Using this formalism, we calculate optimal parameter settings for tracking and subsequent residence time analysis. We apply the theory-suggested parameter set to extract residence times of the transcription factor CDX2 in the cell nucleus from a single molecule experiment ^26^.

## Results

### Single molecule tracking

Our single molecule tracking and analysis framework is designed to simultaneously analyse and compare several multi-movie data sets corresponding to different experimental conditions such as movie acquisition schemes or biochemical treatments, thereby facilitating the scientific workflow (Figure 1 and Supplementary Material). The data-loading tool automatically scans selected folder structures for tiff-formatted movies. Specific regular expressions in filenames can be used to automatically determine frame cycle times or to select for experimental conditions. In addition, accompanying images or movies carrying information about regions of interests (ROIs) such as the cell nucleus can be loaded. We implemented common movie handling and visualization features such as brightness, contrast and z-projection. Movies within the same or a different data set can be conveniently accessed.

**Figure 1.**
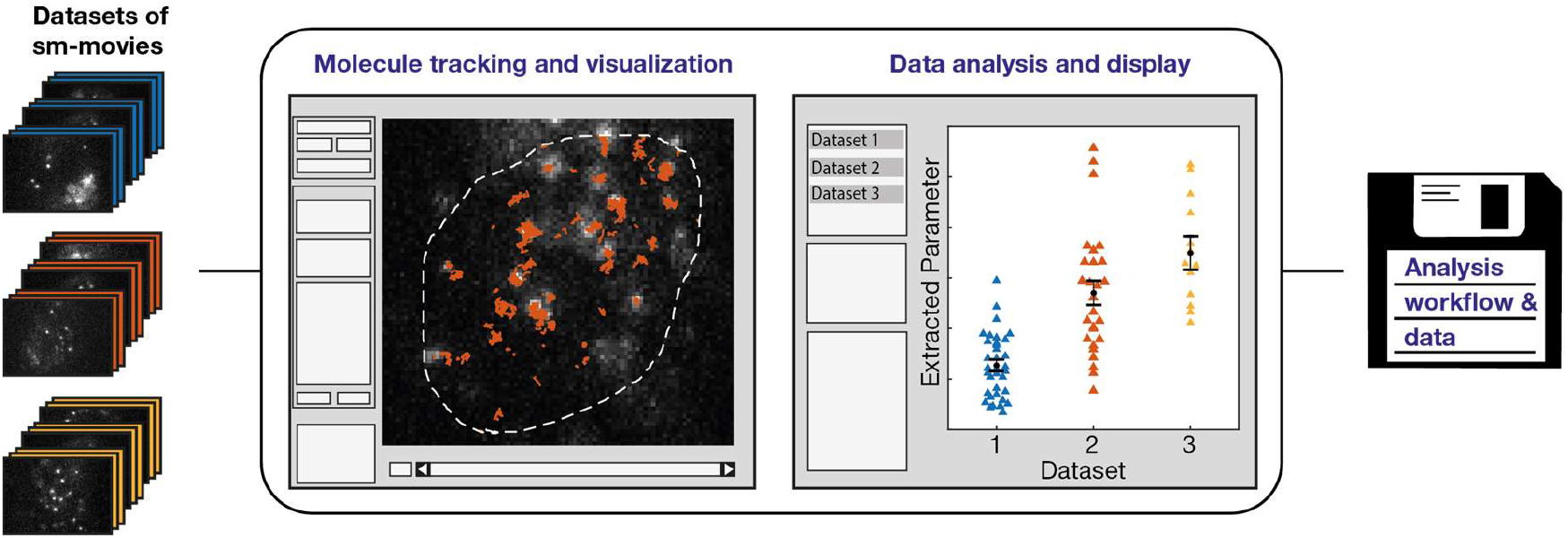
Workflow of TrackIt software. TrackIt enables comparative tracking and analysis of multiple single molecule data sets, each consisting of multiple movies. After loading and manual or automatic determination of tracking parameters, molecules are detected and tracked within a region of interest. Various display options of tracks provide visual feedback on the tracking process. Within the analysis tool, multiple quantitative parameters can be extracted, including bound fractions, residence times, diffusion coefficients, jump angles and confinement radius. Data is stored in a convenient batch structure for later reanalysis or comparison to other data sets.

Our single molecule tracking approach includes four steps to detect individual molecules and link their motion through consecutive images. First, we apply a combination of two wavelet filters to enhance spots representing single molecules ^27^. Wavelet filters performed well in the particle tracking challenge ^2^. Second, we select spot candidates using a local maximum search approach and filtering candidates with a user-defined intensity threshold. Third, we refine the localization of the filtered spots using TrackNTrace’s fast 2D Gaussian fit ^13^.

Finally, we link spots into tracks using a simple model-free nearest neighbour algorithm, which is widely used at low or intermediate spot densities ^2,5,6,12,21^. For high spot densities or a priori known movement models, we implemented u-track as an alternative tracking algorithm ^7^. The nearest neighbour algorithm links spots that are nearest neighbours in two consecutive frames as long as their distance does not exceed a user-defined tracking radius. To account for fluorophore blinking and stochastic fluctuations in spot intensity, the tracking algorithm may bridge missing detections (gap frames) in a user-defined number of subsequent frames ^17^, as long as the first track segment contains a certain number of detections. The choice of tracking radius and concatenating track segments may introduce tracking errors, which we discuss below. Our GUI allows controlling all four steps of the tracking workflow. Importantly, we implemented the possibility to compare different choices of tracking parameters to enable assessing their influence on the tracking results.

Our framework simplifies the effort of analysing numerous movies of multiple data sets by applying the detection and linking steps to all loaded movies without further input by a user. The results and tracking parameters are stored in a single analysis batch structure. This unique data structure summarizes associated files, properties of detected spots and tracks and all tracking parameters in one single file. Thus, reproducing results and comparing multiple processed data sets is possible with minimal effort.

### Data analysis

We implemented a GUI-module to analyse multiple characteristics of single molecule tracks and to display the results, enabling direct comparison of different data sets (Figure 1 and 2, Materials and Methods and Supplementary Material). Tracked molecules can be visualized using a set of intuitive tools allowing both directly inspecting the effect of changes in tracking or analysis parameters and accessing their spatiotemporal dynamics. Besides conventional plotting of spots and tracks, heat maps of localizations 28–30 and jump distances ^31^ can be displayed. We distinguish mobile and immobile molecules based on the time spend within a certain area ^22–24^ (Materials and Methods). Lifetimes of immobile molecules, imaged at different time-lapse conditions to allow for photobleaching correction ^25^, are collected in survival time distributions ^23^. From these, the complete spectrum of dissociation rates is extracted by inverse Laplace transformation using GRID ^26^. Also, the fractions of bound molecules can be assessed using interlaced time-lapse microscopy ^32^. For mobile molecules imaged at sufficient acquisition speed, analysis of the jump distances within each track yields diffusion coefficients ^33^, the bound fraction as amplitude of apparent slow diffusion due to the localization error ^23,33^ and confinement radii as function of the mean displacement ^34^. In addition, histograms of the angles between the jumps of a track can be displayed, informing on compact versus non-compact diffusion ^35,36^. We further implemented the possibility to analyse the intensity profile of tracks ^37^. The batch structure facilitates assessing differences between multiple experimental conditions.

**Figure 2:**
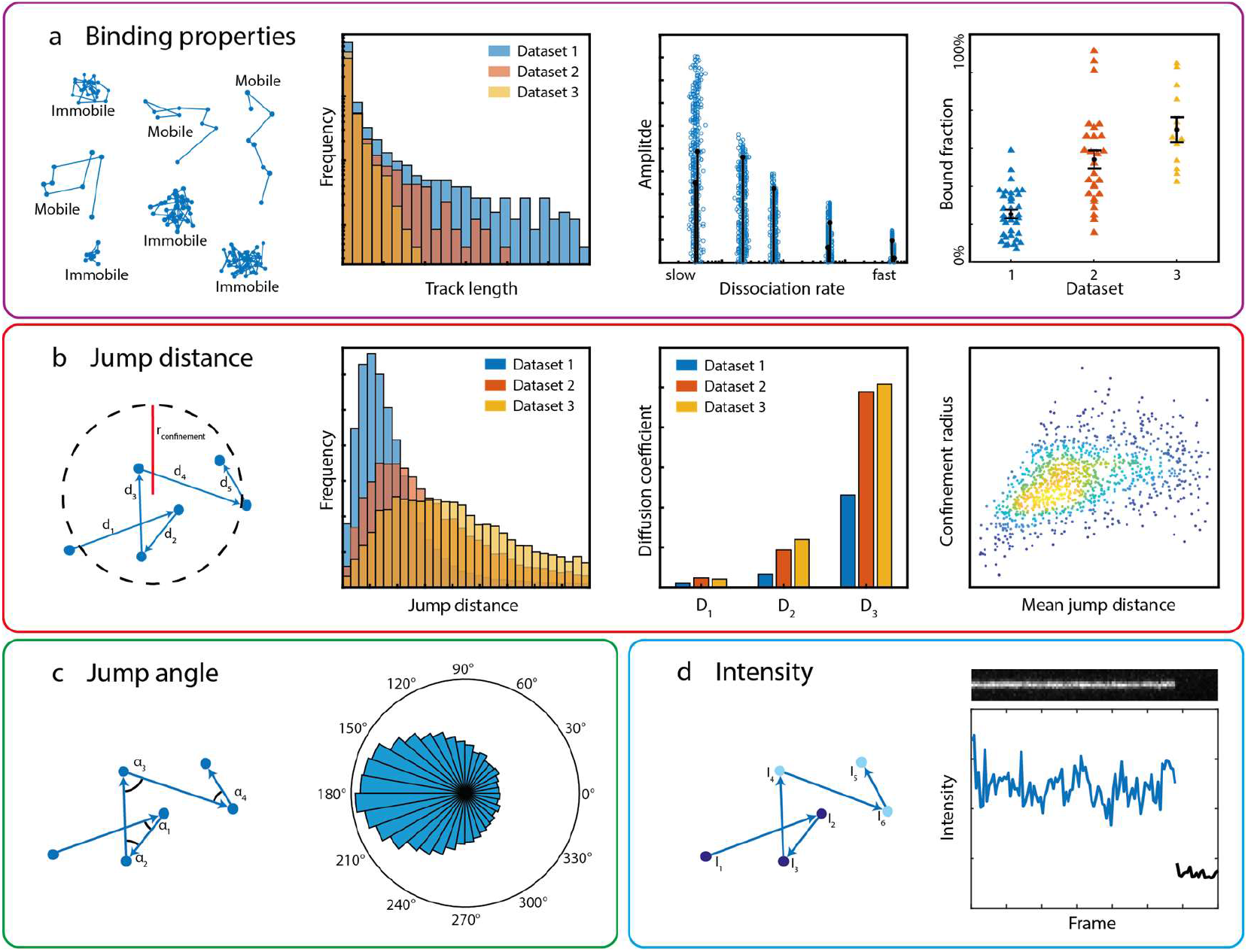
Examples of evaluation parameters and display options. (**a**) For molecules classified as bound by an area threshold, the duration of tracks (track length), the dissociation rate spectrum via GRID and bound fractions can be obtained. (**b**) From the jump distances within tracks, diffusion coefficients and the confinement radius can be computed. (**c**) Angles between consecutive jumps are calculated and displayed in a polar histogram. (**d**) Each track can be individually explored by plotting the intensity over time together with a kymograph.

To ensure transparency and flexibility, we included a data export option, with which track coordinates are stored in specific Matlab or csv file formats to enable utilizing external analysis environments, such as Spot-On or vbSPT for diffusion parameters ^19,20^.

### Description of tracking errors

The track of an immobile molecule stops when (i) the molecule exits the immobile state, (ii) the fluorescent label photobleaches (photobleaching loss) or (iii) a tracking error occurs (tracking loss) (Figure 3a). Only case (i) depends on the properties of the molecule and carries information about its residence time in the immobile state. In contrast, both photobleaching and tracking errors falsely terminate a track and thus blur the survival time distribution if they are not corrected for. Although tracking loss may occur as often as or even exceed photobleaching loss, correction methods for tracking loss are less developed than for photobleaching loss.

**Figure 3:**
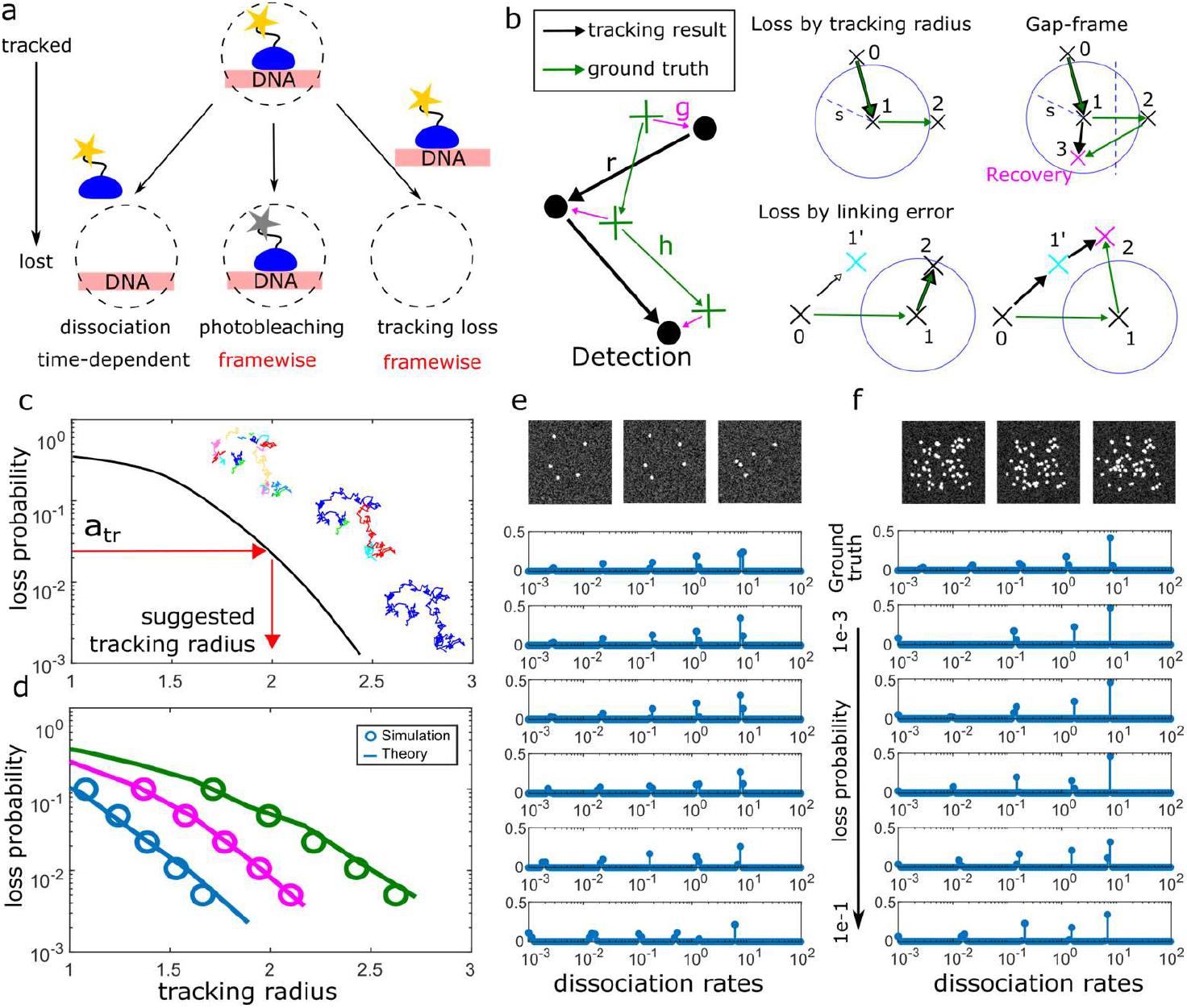
Prediction of loss probabilities in nearest neighbour tracking. (**a**) The track of a bound molecule stops after dissociation, which depends on the laboratory time. Additionally, photobleaching and tracking losses may falsely terminate a track. These losses depend on the frame-cycle time. (**b**) Sketches of tracking losses in the nearest neighbour algorithm. Tracking loss may occur due to jumps out of the tracking radius and due to erroneous links to a different molecule in close proximity. If one gap frame is allowed, both types of losses can be partially recovered. Recovered tracks retain the correct temporal information, however the spatial properties of the track will be altered. (**c**) Simulated loss probability as function of tracking radius. Large tracking radii increase erroneous links to neighbouring detections, small tracking radii increase losses of the tracked molecule. (**d**) Comparison of loss probabilities (spheres) of simulated spots tracked with the nearest neighbour algorithm using different tracking radii with loss probability predicted by theory (lines). Three different time-lapse conditions where simulated, with frame-cycle times of 0.05 s (blue), 1 s (magenta) and 5 s (green). (**e**) and (**f**) Dissociation rate spectra of molecules simulated with five different dissociation rate constants after tracking them using the tracking radii suggested by theory for a given loss probability. (**e**) For low densities, dissociation rate spectra are correctly inferred if the loss probability was below 10% per frame. (**f**) For a density of 0.005 Spots per pixel, only fast dissociation rates were recovered in cases of low loss probabilities.

To be able to account for tracking errors of immobile molecules, e.g. transcription factors bound to chromatin, we established a model for the tracking loss of the nearest neighbour algorithm (Materials and Methods). We considered tracking loss due to jumps out of the tracking radius and erroneous links to a different molecule in close proximity (Figure 3b). If one gap frame is allowed, these losses can be partially recovered and the lifetime of the immobile state stays unaltered (Materials and Methods). Oversize jumps are recovered if the molecule returns back into the tracking radius after the gap frame and is closer to the centre of the tracking area than to its previous position. A falsely linked track can be correctly continued, if the subsequent detection is within the tracking radius of the falsely linked spot. We calculated the probabilities of recovery and final loss by considering the geometry of these situations and obtain an overall probability *a_tr_* for the tracking loss of (Materials and Methods):

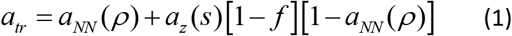

where *a_NN_* is the loss due to erroneous linking, that depends on the spot density *ρ* and *a_z_* is the loss due to jumps out of the tracking radius *s*. The factor (1 − *f*) represents the reduction of the tracking loss by allowing one gap frame. The optimal loss probability balances obtaining fully linked tracks at large tracking radii and low erroneous linking of adjacent molecules (Figure 3c).

To validate our approach, we simulated a single immobile spot without photobleaching and without dissociation from chromatin (Table 1 and Materials and Methods). We linked it with our nearest neighbour algorithm, and obtained the loss probability from the average lifetime of the tracks. We compared the loss probability as function of tracking radius and at different time-lapse conditions with our theoretical prediction (Figure 3d and Table 1). The theoretical expectation well described the *in silico* experiment.

**Table 1:** Simulation parameters for time-lapses data. Scenario 1 corresponds to a single spot that has an infinite residence time and does not exhibit photobleaching. Scenario 2 and 3 correspond to a molecule that exhibits five dissociation rates from chromatin and is subject to photobleaching at different spot densities per frame.

| Scenario | Molecules per frame | Relative frequency | Off | a | Time-lapse conditions | Frames | Point spread function width | Conversion factor pixel to $\mu\text{m}$ |
| --- | --- | --- | --- | --- | --- | --- | --- | --- |
| 1 | 1 | 1 | 1e-15 | 1e-15 | 0.05; 1; 5 | 500 | 1px | 0.16 |
| 2 | 5 | 0.03,0.06,0.130,0.26,0.52 | 3e-3, 2e-2, 0.2, 1.3, 10 | 0.01 | 0.05,0.3,1.5,5 | 400 | 1px | 0.16 |
| 3 | 50 | 0.03,0.06,0.130,0.26,0.52 | 3e-3, 2e-2, 0.2, 1.3, 10 | 0.01 | 0.05,0.3,1.5,5 | 400 | 1px | 0.16 |

### Experimental correction of tracking errors

Tracking errors have a certain probability to occur after each frame of a movie, similar to photobleaching. When measuring the residence time of immobile molecules, tracking errors can be corrected for by applying a time-lapse imaging scheme with several time-lapse conditions, if the loss probability per frame is constant for all time-lapse conditions. In this case, correction using time-lapse imaging is similar to the correction of photobleaching ^25^. The reason is that the residence time of the molecule does not depend on the time-lapse condition, while both tracking errors and photobleaching do. In effect, both photobleaching and tracking errors can be corrected for simultaneously by combining both losses in a single loss probability.

When analysing a time-lapse experiment, a different tracking radius has to be chosen for each time-lapse condition to yield an overall constant loss probability (Figure 3d). We implemented an algorithm in our tracking and analysis framework, which, for a user-defined loss probability, calculates the corresponding tracking radius for each time-lapse condition.

We tested the performance of our correction approach in the analysis of residence times of immobile molecules. We simulated a time-lapse experiment for a scenario where immobile molecules at a low density of 5 spots in 100 × 100 px were subject to five different binding interactions with corresponding dissociation rates (Figure 3e, Table 1 and Materials and Methods). We tracked the molecules by specifying a loss probability and using the suggested tracking parameters for each time-lapse condition in the nearest neighbour algorithm. Subsequently, we used GRID to extract the spectrum of dissociation rates ^26^. To determine to which extend tracking errors influenced the result we varied the loss probability over two orders of magnitude. We found that for high loss probabilities > 10%, small dissociation rates arising from long-lasting tracks could not be inferred correctly. In contrast, for low loss probabilities < 10%, the ground truth was well inferred with our analysis.

We further tested to which extend the density of molecules affected our approach. We simulated densities up to 50 spots in 100 × 100 px (Figure 3f and Table 1) in steps of 0.0005 spots per pixel to provoke incorrect linking between individual spots. Molecules were again subject to five different binding interactions. The maximal density of molecules at which dissociation rate spectra and thus associated residence times could be well inferred was 0.0025 spots per pixel.

### Analysis of the dissociation rate spectrum of the transcription factor CDX2

Finally, we applied our tracking and analysis framework and the algorithm which predicts optimal tracking parameters to re-analyse the dissociation rate spectrum of the transcription factor CDX2 using previously published single molecule imaging data (Figure 4) ^26^. The data consists of four time-lapse microscopy conditions of a HaloTag-CDX2 fusion protein labelled with a SiR dye. Time-lapse movies were obtained with 50 ms frame acquisition time and an overall frame cycle time of 0.05 s, 1 s, 5 s and 9 s. In our tracking parameter prediction algorithm, we set the loss probability to 0.01 and the algorithm calculated a corresponding set of tracking parameters for each time-lapse condition. As a result, the algorithm revealed tracking radii and the corresponding time periods during which a HaloTag-CDX2 molecule should stay within this boundary to be identified as bound. The tracking radii where 100 nm, 240 nm, 410 nm and 430 nm for the frame cycle time condition of 0.05 s, 1 s, 5 s and 9 s, and the minimum track length was 5 frames for 0.05 s frame cycle time movies and 2 frames for the other conditions. For tracking, we used the nearest neighbour algorithm and allowed bridging detection gaps of one frame as long as a track already existed for at least 2 frames. Next, the resulting track durations were transferred to the GRID toolbox to extract the dissociation rate spectrum. Our approach with computationally determined tracking radii and the previous analysis using manually chosen tracking radii yielded comparable dissociation rate spectra (Table 2). The main deviations are in the amplitudes, not the values of dissociation rate clusters. For future experiments, computationally determined optimized tracking radii will ensure robust data analysis.

**Figure 4:**
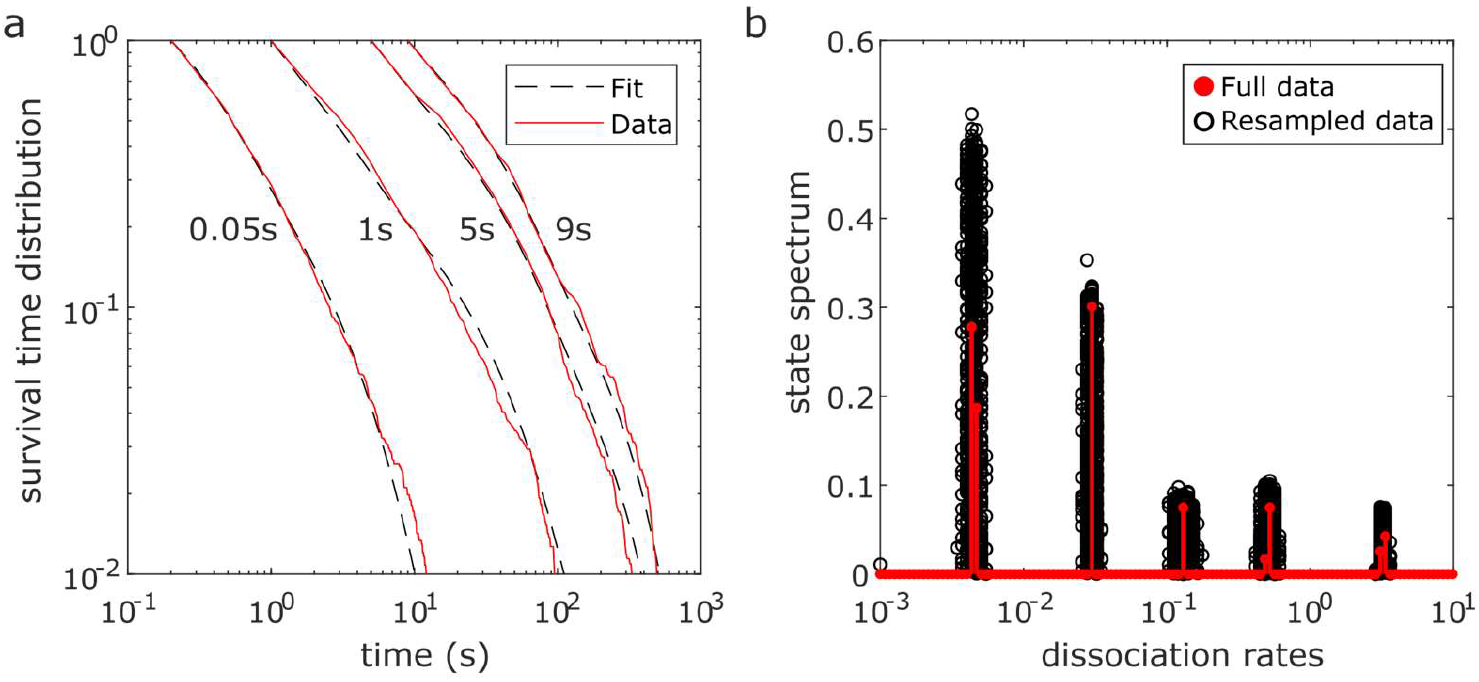
Dissociation rate spectrum of CDX2 – chromatin interactions. Live-cell single molecule movies of SiR-Halo-CDX2 ^26^ where analysed using the tracking radii suggested by theory for a tracking loss probability of 1e-2. (**a**) Fluorescence survival time distributions of tracked molecules (red lines) at different time-lapse conditions (indicated on top) and survival time functions obtained by GRID (black dotted lines). (**b**) State spectrum of SiR-Halo-CDX2 obtained by GRID using all data (red circles) and a superposition of 499 GRID results obtained by resampling 80% of data (black circles) as an error estimate of the state spectrum.

**Table 2:**
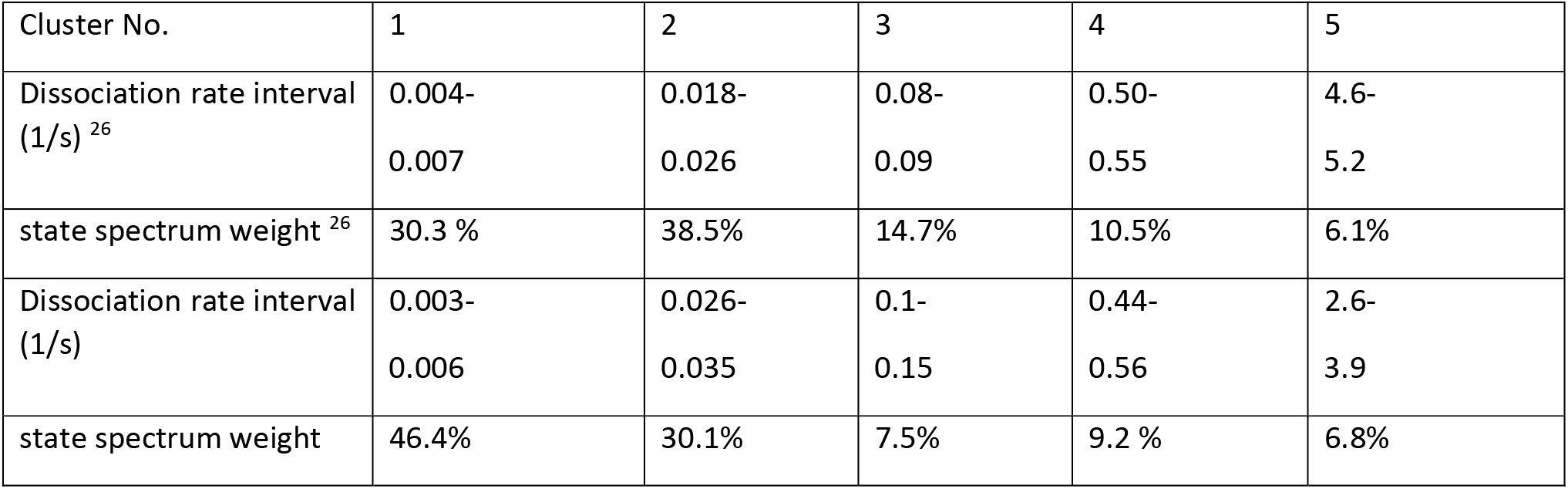
Comparison of the inferred CDX2 spectrum with ^26^. Dissociation rate interval specifies the manually assigned dissociation rate intervals corresponding to a dissociation rate cluster. The spectral weight of each of the five distinct clusters of the CDX2-spectrum was obtained by integrating the GRID amplitudes resulting from 100% of the measured survival times.

## Discussion

We introduced the tracking and analysis framework TrackIt, which simplifies analysing and comparing multiple different large single molecule fluorescence data sets due to a comprehensive list of GUI modules and a batch structure. Thus, fast and reproducible analysis can be performed without the need for additional manual steps or programming. In particular, TrackIt is well suited to systematically compare different settings of tracking parameters and to scan data sets differing in experimental conditions for differences in kinetic or structural parameters.

We further introduced quantitative considerations of tracking losses of immobile molecules in a nearest neighbour tracking algorithm and implemented means to partially correct for them. The nearest neighbour algorithm is more prone to linking errors than more complex algorithms ^2,7,8^. However, in contrast to complex approaches that produce unpredictable tracking losses, the nearest neighbour algorithm allows for a theoretical prediction of tracking losses. This prediction allowed us to automatically determine consistent settings of tracking parameters for the analysis of immobile molecules. We note that tracking errors in every tracking algorithm can be minimized by measuring at low molecule densities, thereby trading optimized tracking for measurement throughput. Overall, we provide a pipeline to analyse transcription factor residence times including both photobleaching and tracking error corrections.

Each research question comes with own requirements for data analysis. Thus, while our framework provides state-of-the art analysis approaches applied in recent publications, care has to be taken whether an implemented approach optimally covers the analysis needs. For example, analysing bound fractions requires separating molecules into different kinetic classes. While we implemented a commonly used approach to identify bound molecules by their restricted mobility area ^22–24^, alternative classification approaches have been published ^34,38–40^. Moreover, the bound fraction analysis we implemented is best suited for data sets captured using interlaced time-lapse microscopy ^32^. Some important analysis approaches are not yet implemented in TrackIt, for example the possibility to analyse two-colour single molecule data. However, being implemented in commonly used Matlab format, our framework constitutes a broadly accessible platform to which novel analysis schemes can be added.

We used theory-suggested tracking parameters and GRID to determine the spectrum of dissociation rates of CDX2 proteins. We could verify our previous results, where manually assigned tracking parameters were carefully adjusted considering movies of all time-lapse conditions. Our theory-predicted tracking radii reach the same quality as manual parameters and thus enable reproducible, user-independent data analysis.

## Materials and Methods

### Localization and jump distance mapping

The spatial distribution of localizations and mobility parameters of tracked molecules contains valuable information about their function and environment ^28,30,31^. We enabled creating a super-resolved heat map of localizations by using all detected spots whose position is determined to sub-pixel precision with a 2D Gaussian fit. The positions of all spots were then accumulated in a 2D histogram. The pixel values therefore correspond to the amount of detections in each pixel. The bin size of the histogram can be chosen as a multiple of the original pixel size, which results in an up- or downscaled image. Similarly, we create a heat map of jump distances. We define a jump as a change in position between two consecutive localizations of a track. For all jumps within a track, a virtual line is drawn between the start and end positions of a jump. Each pixel touching this line is assigned with the corresponding jump distance. The resulting 2D histogram is then normalized by the amount of jump events in each pixel. Again, the image can be up- or downscaled by using an appropriate bin size of the histogram.

### Jump distances and diffusion Analysis

To obtain a direct overview of the jump distance distribution of all tracks, a histogram is created using all single molecule jump distances between consecutive frames. The bin size Δ*r* of the histogram can be set individually. In addition, the number of jumps of a tracked molecule to be considered in the histogram can be chosen. If only one jump per track is chosen, a biased weight of immobile over mobile molecules is minimized.

To extract diffusion coefficients from mobile molecules, we created a cumulative histogram of the squared jump distances, normalized to the total number of jump events. We fitted the resulting cumulative density distribution of squared displacements with a Brownian diffusion model including either two or three different diffusion components ^23,25,33^. For the two-component model we used

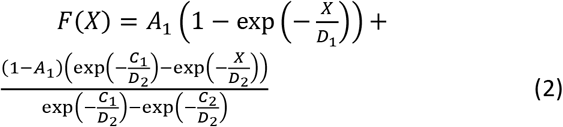

while for the three-component model we used

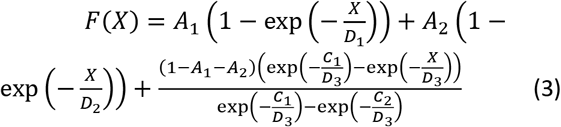

Here, *D_i_* denotes the diffusion constant of the i-th component, *X* = (*x*^2^ + *y*^2^)/(4*τ*) with the camera frame cycle time *τ* and *A_i_* denotes amplitude. The last term is normalized by 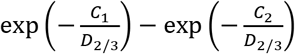 to account for the cut off due to the lower limit of jump distances, which is C_1_ = 0, and the upper limit of jump distances, C_2_, which is given by the tracking radius, respectively.

Using the GUI, the original cumulative histogram is overlaid with the fitted function for visual inspection. Moreover, the resulting diffusion constants and amplitudes are displayed and compared between different data sets. We give 95% confidence intervals as an estimate of the error of fit-parameters. Adjusted R^2^ values are used to estimate how well the data is represented by one of the models.

### Confinement radius analysis

We implemented a two-parameter representation of tracks in which tracks are sorted according to their confinement radius and their mean jump distance ^34^. This representation gives insights into different mobility classes of single molecules. We calculated the confinement radii as previously described ^34^. In brief, the mean squared displacement as function of time of each track is fitted with a confined diffusion model ^12,34^

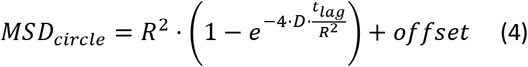

where R is the radius of confinement, D is the local diffusion coefficient, and the offset is introduced to account for the finite localization precision. In order to select only confined tracks for analysis, the mean squared displacement as function of time of each track is fitted with a power law *MSD* = 4 · *D* · *t^α^*, where *D* is the diffusion coefficient and α an exponent that indicates the motion type ^34^. A threshold can be chosen for the maximum value of α that should be considered for further analysis.

### Analysis of the dissociation rate spectrum and corresponding residence times

For immobile molecules, we implemented the possibility to analyse their dissociation rate spectrum and corresponding residence times. To obtain the dissociation rate spectrum, the survival time distribution of track durations of all tracks in continuous video or a time-lapse data set were calculated in the GRID toolbox. In time-lapse microscopy, the acquired frames are separated by a time period without illumination of the sample. Next, dissociation rate spectra were extracted in a global analysis of all time-lapse data sets using GRID ^26^. In brief, solving the inverse Laplace transformation of each survival time distribution is translated into a single minimization problem that can be handled by a gradient method.

### Bound fraction analysis

The fraction of molecules bound to a stable structure such as chromatin can be determined by interpreting the amplitudes of diffusion components of tracked molecules ^23,33^. In another approach, the fractions of molecules belonging to two different binding time classes can be approximated using the interlaced time-lapse microscopy (ITM) illumination scheme ^32,41^. In ITM, two subsequent frame acquisitions are followed by a longer dark time. Detected molecules are sorted into different binding time classes. Tracks, which survive at least one dark period, are classified as long bound, tracks which persist for two consecutive frames are classified as short bound and single detections are classified as unbound/diffusing. To obtain accurate fractions, they have to be corrected for photobleaching ^32^. Continuous movies may contain information comparable to ITM, however will be highly affected by photobleaching.

Ones classified, we calculate the overall bound fraction using

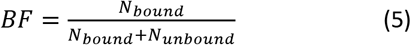

Where *N_unbound_* denotes the number of bound molecules and *N_unbound_* the amount of unbound molecules.

In order to distinguish transient and stable bound molecules we calculate the fraction of long bound molecules

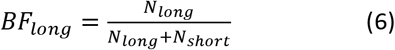

Where *N_long_* denotes the number of long bound molecules and *N_short_* the amount of short bound molecules. A determination of bound fractions for each movie leads to a variation in the bound fraction from movie-to-movie and the final bound fraction is calculated from the mean value over all movies. To avoid an over-representation of movies with low molecule counts, events for each binding time class are additionally summed over all movies resulting in a single “pooled” bound fraction of all movies.

### Intensity and kymograph analysis

We implemented the possibility to select individual trajectories and plot them in a separate window together with its intensity and the associated kymograph ^37^. To visualize the intensity over time, the mean intensity in a 3×3 pixel window around the spot centre is calculated and plotted until the track is lost. Kymographs are visualized in separate plots for each spatial dimension. One spatial dimension is displayed versus time, while the other spatial dimension is maximum projected. After the end of track, both intensity plot and kymographs are continued for the next 20 frames. This allows controlling the quality of the tracking process and assessing whether the molecule photobleached in a single or in multiple steps.

### Jump angle analysis

To analyse the angles between consecutive jumps ^35,36^, we calculated the scalar product between the two normalized vectors representing the directions of the two consecutive jumps. We then took the inverse cosine to calculate the angle between the two vectors. The resulting angles are visualized in an angular histogram ^35^.

### Simulation of single molecule time lapse data

We generated single molecule movies using a custom simulator implemented in Matlab 2019a. We simulated diffusion of a protein and photobleaching of an attached fluorescent label. Diffusion is altered upon association to or dissociation from chromatin by the protein. The protein can enter different binding states that are characterized by their on- and off-rates.

In favour of fast simulations, we renounced simulating the molecule position at fixed small time steps, but used the Gillespie direct method. With this approach, motion blur is not included in the simulation. We first determined the times until a photobleaching event or until a transition event from a diffusing to a bound state or vice versa. We then determined the position of the molecule for each frame. In case a state transition occurred outside a frame interval, we also determined the position of the molecule at the corresponding time. Jump distances were drawn from the 2D diffusion probability density corresponding to the diffusion coefficient D of the current state (free diffusion or apparent diffusion due to the localization error if bound).

We then used the simulated trajectories to generate images. For each spot, we simulated a point spread function with intensities corresponding to a lognormal photon count distribution. To add background, we used uniform random numbers for background noise. To approximate non-uniform background, we applied a band-pass filter, which additionally enhanced features of the background.

### Tracking loss in nearest neighbour tracking

The nearest neighbour algorithm links detected spots into tracks by comparing their positions in two consecutive frames. It links spots that are closest to each other if their distance does not exceed a certain tracking radius. Spots that cannot be linked to an existing track start a new track. Errors occur if a jump distance is larger than the tracking radius or if the track is erroneously linked to a different molecule in close proximity.

To estimate the probability of losing a track we assumed that the detected position of a spot depends on diffusion and a localization error. The effective squared jump distance between two consecutive frames is given by

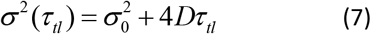

which results in the corresponding probability density for the detected position

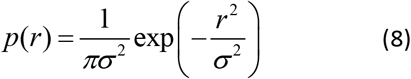

The probability *a_tr_* of the spot position to exceed the tracking radius *s* is obtained by integrating the above equation over the interval 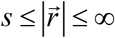

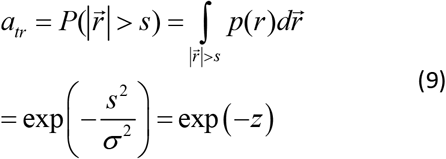

where we introduced the dimensionless variable

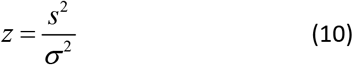

We further considered an interruption of the track due to an erroneously linked molecule. The position of the disturbing molecule is denoted by 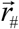. In order to disrupt the track, the molecule has to be closer to the second detected position. The probability to lose a track due to linking errors in dense environments is obtained by considering all possible configurations of this scenario:

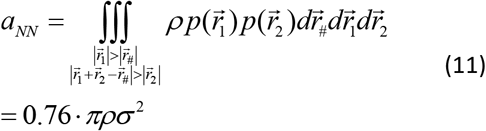

where *ρ* is the density of spots/frame.

### Partial recovery of tracking loss by introducing one gap frame

Tracking loss can be partially recovered if the tracking algorithm allows bridging one frame with missed detection.

We first calculated the probability *I* that two consecutive jumps including localization error lie within a given area Ω:

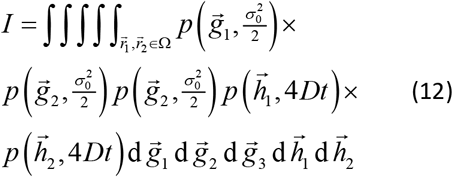

with the integration borders

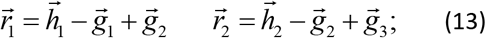

where h is the jump of the molecule and g is the detected position. By inserting the probabilities (12) and detected spot positions (17) in (16) and integrating with respect to 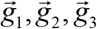 we obtain

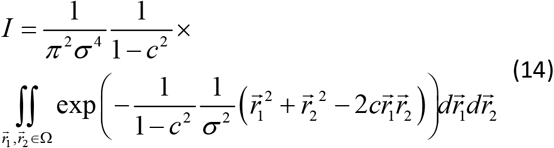

We next used *I* to estimate the effect of a gap frame. We calculated the probability *I* (Ω) of a molecule to return inside the tracking radius after having left it in the previous frame. For this case, we need to consider that the molecule outside the tracking radius was detected but not linked to the existing track and therefore started a new track. The detection in the current frame has to be closer to the starting position than to the molecule outside the tracking radius otherwise the track will be cut. The area Ω_*DF*_ corresponding to this situation is given by:

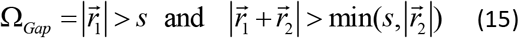

Solving the Integral (18) with area Ω_*Gap*_ yields the probability that the track is recovered. The integral *I* (Ω) was calculated numerically. The corresponding probability to lose a track after a gap frame is given by 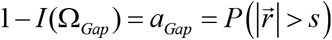. Compared with the tracking loss without gap frame, the probability to lose a track if tracked with a gap frame is upmost a factor of 0.5 smaller.

The overall probability *a_tr_* for losing a track in presence of a gap frame and in presence of erroneous linking is given by:

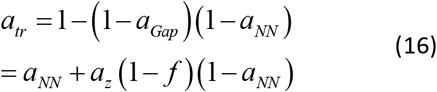

### Calculation of tracking radius to ensure a certain loss probability

To obtain the tracking radius for a given loss probability we need to solve equations (13) and (7) for *s*. Since no closed-form equation can be given for *s*(*a_tr_*), we employed an iteration scheme. We started the iteration at a starting point *s*_0_ and iterated until the change between the i-th and i+1-th iteration is smaller than 1e^−2^. The equations for the step *i* → *i* + 1 are given by

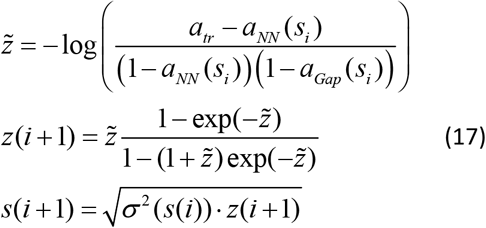

In our iteration, we accounted for the fact that jump distance distributions are cut by the tracking radius ^12^. In each iteration, we determined the mean of squared jump distances *σ*^2^(*s_i_*) by tracking with the nearest neighbour algorithm with tracking radius *s_i_*.

### Simulation Parameters

We simulated videos with frame sizes of 100 × 100 pixels. The SNR of spots was chosen as 25. The diffusion constant of chromatin bound molecules was set to D=1e-3 μm^2^ s^−1^, the free diffusion constant was set to D=10 μm^2^ s^−1^

## Supporting information

TrackIt manual

## Acknowledgements

We thank members of the Gebhardt and Michaelis labs for helpful discussions. The work was funded by the European Research Council (ERC) under the European Union’s Horizon 2020 Research and Innovation Program (No. 637987 ChromArch to J.C.M.G.) and the German Research Foundation (SPP 2202 GE 2631/2–1 and GE 2631/3–1 to J.C.M.G.). Support by the Collaborative Research Centre 1279 (DFG No. 316249678) is acknowledged.

## Contributions

T.K., J.H. and J.C.M.G. designed the project; T.K. programmed TrackIt; J.H. modelled tracking errors and contributed to programming TrackIt; T.K. and J.H. analysed CDX2 data; T.K., J.H. and J.C.M.G wrote the manuscript.

## Data availability

Data supporting the findings of this manuscript will be available from the corresponding author after publication upon reasonable request. Single particle movies of CDX2, described in ^26^, and tracking results will be made available upon publication.

## Code availability

The TrackIt software is freely available. A Matlab version of TrackIt is available at https://gitlab.com/GebhardtLab/TrackIt.

