## Supplementary material for "Single molecule tracking and analysis framework including theory-predicted parameter settings": TrackIt manual

Release 1.0

written by

**Gebhardt lab, University of Ulm**

Nov. 2020

### 1 General Information

#### 1.1 Licence

This program is free software: you can redistribute it and/or modify it under the terms of the GNU General Public License as published by the Free Software Foundation, either version 3 of the License, or (at your option) any later version. This program is distributed in the hope that it will be useful, but WITHOUT ANY WARRANTY; without even the implied warranty of MERCHANTABILITY or FITNESS FOR A PARTICULAR PURPOSE. See the GNU General Public License for more details. You should have received a copy of the GNU General Public License along with this program. If not, see `http://www.gnu.org/licenses/`.

#### 1.2 Requirements

- Operating system: Windows 10 64-bit
- Matlab version 2019a and above
- For full functionality, the following Matlab toolboxes are required: Optimization, Image Processing, Statistics and Machine Learning, Parallel Computing
- GRID toolbox for analysis of dissociation rates and for prediction of tracking radii `https://gitlab.com/GebhardtLab/GRID`

#### 1.3 Installation

1. TrackIt is available from `https://gitlab.com/GebhardtLab/TrackIt`
2. Extract
3. Install the GRID toolbox ("GRID_for_trackit.mltbx") that comes with TrackIt.
4. Open the TrackIt_v1_0.m file with Matlab
5. Run TrackIt by clicking the run button (F5) or from the command line

### 2 Workflow

#### 2.1 Overview of the main GUI

TrackIt is a tracking and analysis pipeline for single-molecule fluorescence microscopy movies. The Software uses a unique batch data structure to handle and save all data created by the user, it can be stored in a .mat file and loaded into the software at any point of the workflow. Only one .mat file is needed per data-set which makes it easy to analyze and compare many sets of experiments.

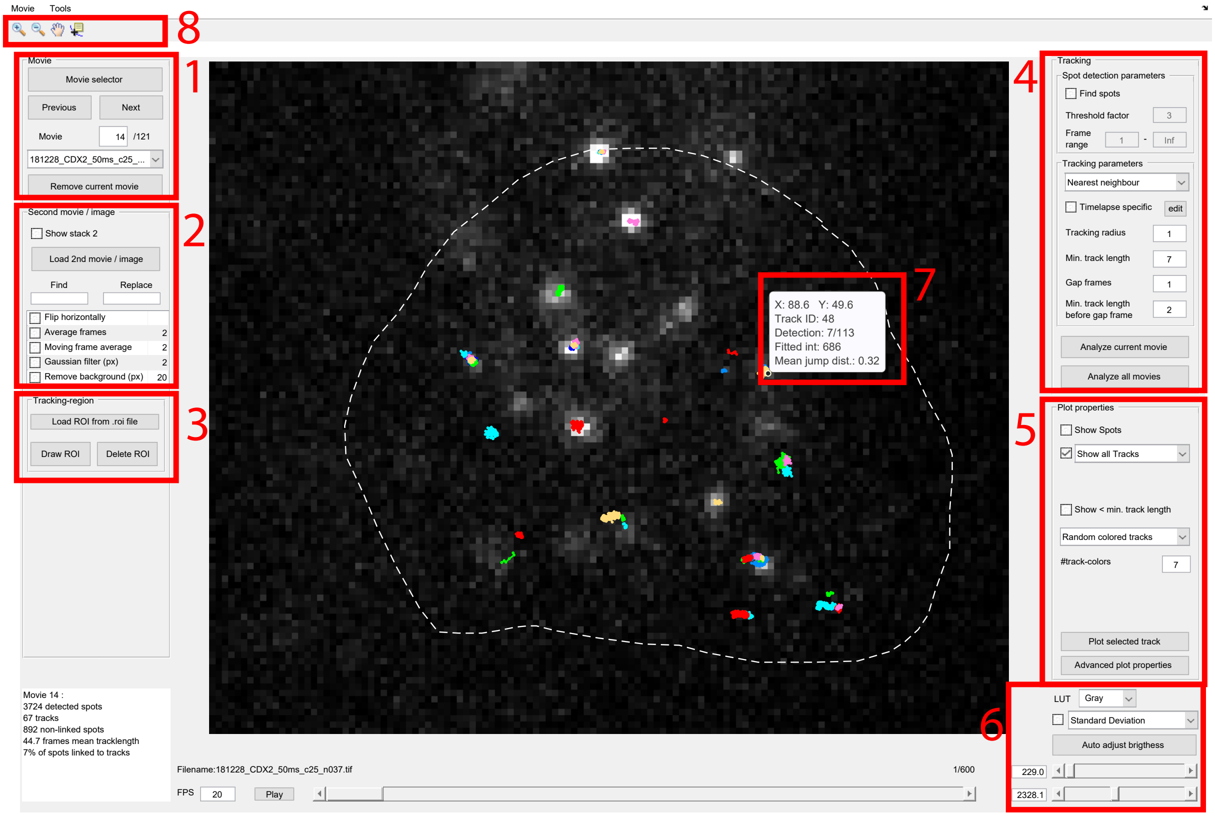

Figure 2.1: Screenshot of the main user interface. 1) Movie panel 2) Second movie / image panel 3) Tracking-region panel 4) Tracking panel 5) Plot properties panel 6) Z-projection and brightness adjustment 7) Toolbar 8) Information on selected track.

1. **Movie panel** Add movies via the "Movie selector" button. Navigate through movies of the current batch with the "next" and "previous" button, by entering a movie number or by selecting a movie from the drop-down list. Single movies can be removed from the current batch with the "Remove current movie" button (the movie is not deleted from your folder).

2. **Second movie/ image panel** Load reference channels carrying information about regions of interest (ROIs) such as the cell nucleus via the "Load 2nd movie / image" button. "Replace" and "Insert" fields help selecting the filename directly without the need of searching through file lists. The reference movie or image can be manipulated with common filtering and averaging operations.

3. **Tracking-region panel** To restrict tracking to a specific region, e.g. the nucleus, a ROI can be drawn or loaded from an existing .roi file which is created automatically every time a ROI is drawn.

4. **Tracking panel** All relevant parameters concerning spot detection and tracking can be set. By clicking "Analyze current movie" or "Analyze all movies" fluorescent molecules are tracked in the current movie or all movies, respectively. To analyze movies with several tracking parameters at once, comma separated values can be entered in the parameter fields. Tracking parameters can also be applied with respect to the corresponding frame-cycle-time of each movie by checking the "Timelase specific" box.

5. **Plot properties panel** All plot related properties such as size, style and coloring of spots and tracks, the range in which tracks are shown and scalebar options are accessible here. Clicking the button "Plot selected track" opens a window showing kymograph, intensity and position plot of a track (see Fig. 2.6) selected with the "Data tip" in the toolbar .

6. **Z-projection and brightness adjustment** Settings concerning displayed pixels as the color lookup table or brightness and contrast can be adjusted in the lower right corner. Additionally z-projections of the displayed movie or detection and jump distance mappings can be displayed.

7. **Toolbar** Toolbar with tools to zoom in and out, move the plot area and to select specific tracks (see below).

8. **Information on selected track** Selecting a track with the "Data tip" in the toolbar shows basic track information such as track number, current position, fitted intensity and mean jump distance.

#### 2.2 Import movies using movie selector

Clicking the "Movie Selector" Button in the main GUI opens the "Movie selector" window.

Movies can be added in two ways:

1. **Add movies button** .tiff files can be selected directly.
2. **Search folder button** All .tiff files in a selected folder will be added. Optionally, a search string can be entered before clicking "search folder", so that only filenames containing the user specified string are added.

**Automatic recognition of frame cycle times**
The software is able to recognize the frame cycle times (the time between two consecutive frames) if the filename contains a string consisting of an underline followed by a number and either "ms" (milliseconds), "s" (seconds) or "Hz" (Hertz). For example "_50ms", "_1s" or "_50Hz" (see Fig 2.2). The files will then be grouped according to their frame cycle time. The number of files of each frame cycle time will be displayed together with the time in milliseconds.

**Manually set frame cycle times**
If no frame cycle time is recognized, the corresponding field will display -1. The time can be manually entered by clicking the frame cycle time field. Movies of different timelapse conditions have to be added separately while entering the timelapse condition for each case.

**Set movie order**
The user can choose on how the movies should be sorted:

1. Filename: Movies are sorted by their filename
2. Frame cycle time: Movies are sorted according to their frame cycle time (or timelapse condition)
3. String pattern: Movies are sorted by the part of the filename that follows a given string pattern (eg. "_n"), ignoring the part of the filename before the string pattern.

By clicking "OK" the movie selector closes and the first movie will be loaded and displayed in the main GUI. Now is a good time to save the workflow for the first time by clicking "File" » "Save batch file as".

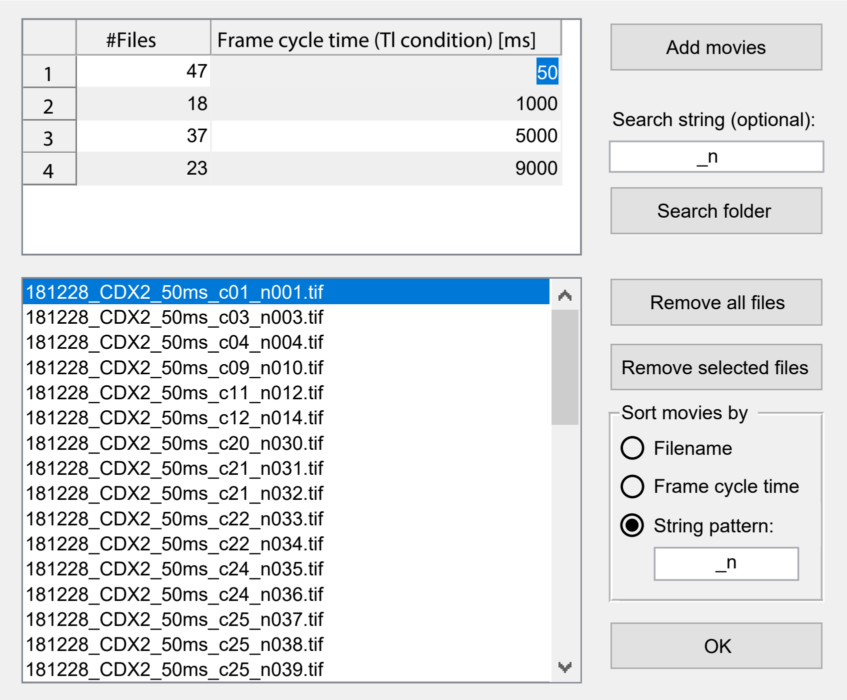

Figure 2.2: Screenshot of transcription factor CDX2 data-set displayed in the movie selector. The search string "_n" was used in combination with the search folder function to add all files in a folder containing that string. The software automatically recognized the frame cycle times (timelapse conditions) as given by the filename.

#### 2.3 Load reference channel

The tracking channel can be overlaid with a second .tiff movie or image (e.g. brightfield image, DAPI or membrane marker). This can be useful for drawing a region of interest (ROI) or localization of proteins with respect to cellular compartments.
A reference channel can be loaded by pressing "Load 2nd movie / image".

In order to ease a laborious search through lists of filenames, a replacement function is implemented. This can be used if the tracking movie and the reference channel share a similar filename for example "cell1_488nm_n001.tif" and "cell1_561nm_n001.tif" and the same folder. The string to replace must be written in the field depicted with "Find", likewise the string which should be inserted must be written in the field depicted with "Replace" (see Fig 2.3). When "Load 2nd movie / image" is pressed the desired filename is directly inserted in the file selection dialog and can easily be opened by hitting "Open".

For the reference channel, following post-processing steps can be applied:

- Flip horizontally: Useful for two color-experiments where a second detection path is split-of by a beam-splitter.

- Average frames: The second movie is grouped and averaged in frame packages given by a user specified number. The amount of frames is reduced by a factor equal to this number
- Moving frame average: The second movie is averaged by calculating a moving average of the frames. Each frame is built by averaging over a user specified amount of frames before and after each frame. The amount of frames therefore stays the same. The window size is automatically truncated at the beginning and the end of the movie.
- Gaussian filter: Uses Matlabs "imgaussfilt" function to apply a 2-D Gaussian image smoothing filter with a standard deviation (width of the filter kernel) specified by the user (in pixels).
- Remove Background: Subtracts a morphologically opened image from the original image. The radius of the disk element used for the opening operation can be specified by the user (in pixels).

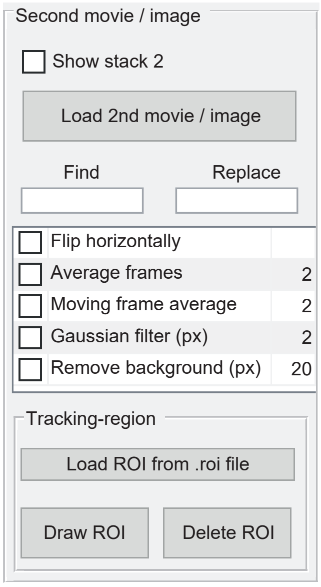

Figure 2.3: Screenshot of the reference channel and ROI panel.

#### 2.4 Region of interest

If desired, a region of interest (ROI) can be drawn. Spot detection will then be restricted to this specific area. If no ROI is drawn, the whole image will be used for detection and tracking.

Directly after drawing is finished, the software will save the ROI in a separate .roi file in a folder called "ROI" located in the same folder as the original movie. Whenever movies are selected via the "Movie selector", the software will check whether a .roi file exists for this file and will load it automatically. .roi files can also be loaded by clicking "Load ROI from .roi file".

#### 2.5 Tracking

Before tracking is started, several parameters can be adjusted in the "Tracking" panel (see Fig. 2.4).

##### 2.5.1 Spot detection and tracking parameters

**Threshold factor** A user defined threshold factor is used to calculate an automatic threshold for spot detection. The threshold is calculated for each movie separately providing comparable detection thresholds between all movies. Reliable detection of single fluorophores is commonly achieved with values between 1 and 5. Movies with a good single-molecule signal (i.e. high signal-to-noise ratio) can usually be analyzed with higher threshold factors.

**Frame range** Specifies the frames between which the spots should be detected and tracked

**Tracking algorithm** Ideally the concentration of fluorescent molecules is low enough so that each spot is spatially well separated from others at all times. In this case the nearest-neighbor delivers fast and reliable tracking results. For higher concentrations the u-Track algorithm might provide better results. For spot density considerations see section 2.7.4, Avg. #spots per frame.

**Tracking radius** Maximum distance at which spots are linked between two consecutive timepoints. In case of several different frame cycle times as used eg. in time-lapse experiments, consistent tracking radii can be predicted automatically, see 2.5.3.

**Min. track length** Minimum number of frames a fluorescent molecule has to persist to be accepted as a track.

**Gap frames** Number of frames a fluorescent molecule can disappear or stay undetected so that tracking is still continued and the fluorescent molecule is combined into a single track.

**Min. track length before gap frame** Number of frames a track has to exist before closing of gaps (gap frames) is allowed. This can help to prevent connecting random detections in overly crowded movies or movies with bad signal-to-noise ratio (SNR).

**Timelapse specific tracking parameters** Increasing movement of cellular compartments (eg. chromatin diffusion) with longer frame cycle times, makes it necessary to chose less restrictive tracking parameters. Therefore, above described tracking parameters can be set dependent on their frame cycle time (timelapse condition) (see Fig 2.5).

In some cases it might be desired to try out several combinations of tracking parameters. This can be achieved by inserting comma separated values into the fields of above described parameters. The software will run through all combinations of tracking parameters and save one batch file per parameter set to a folder specified by the user.

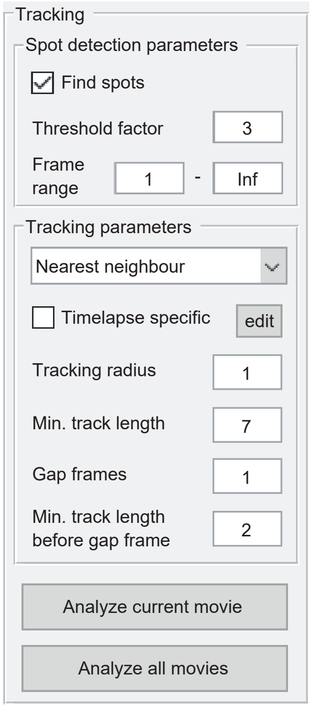

Figure 2.4: Screenshot spot detection and tracking parameter panel.

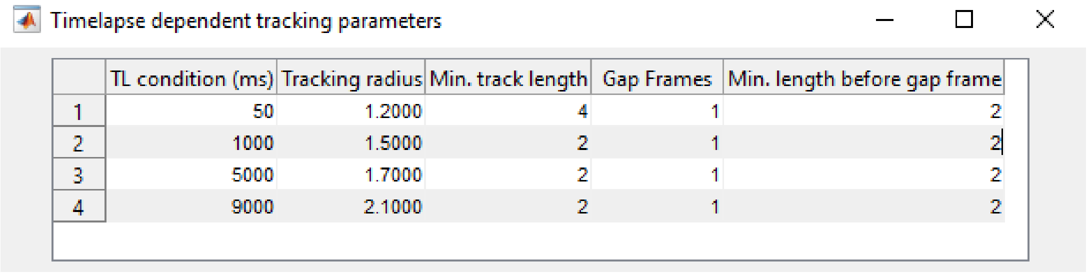

| | TL condition (ms) | Tracking radius | Min. track length | Gap Frames | Min. length before gap frame |
|---|---|---|---|---|---|
| 1 | 50 | 1.2000 | 4 | 1 | 2 |
| 2 | 1000 | 1.5000 | 2 | 1 | 2 |
| 3 | 5000 | 1.7000 | 2 | 1 | 2 |
| 4 | 9000 | 2.1000 | 2 | 1 | 2 |

Figure 2.5: Screenshot of the timelapse specific tracking parameters window where a different set of tracking parameters is used for each of the four timelapse conditions.

##### 2.5.2 Tracking routine

By clicking either "Analyze current movie" or "Analyze all movies" tracking is performed with the predefined parameters. The tracking routine consists of four steps:

1. **Filtering raw data** Movies are filtered using a wavelet filter as described in [5].
2. **Detecting candidates** First a threshold is applied to the wavelet filtered image to separate signal from background noise. The threshold is calculated by multiplying a user defined threshold factor with the standard deviation of the background noise. Candidate detection is based on local maxima finding at the pixel level. In brief, all pixels that have the same value before and after image dilation are selected.
3. **Position refinement** Candidate positions are refined to a sub-pixel precision by fitting a Gaussian function using the freely available psfFit_Image.m from the TrackNTrace software as described in [12]. For spots laying closer together than 2 pixels, the smaller peak is discarded after fitting in order to avoid multiple detections per spot.
4. **Tracking** Linking spots in time and space can be carried out using either a fast nearest-neighbor algorithm or the more sophisticated u-Track algorithm of [7].

##### 2.5.3 Automatic determination of tracking parameters

Click "Tools" » "Predict tracking radii" to open a separate window for tracking radii prediction. The table gives an overview of all parameters. Before running the program, choose wheter gap-frames shall be allowed (=1) or not (=0). Furthermore the shortest track length can be adapted. Continue by clicking "execute". Fill in the inverse of the trageted track length, i.e. for a targeted mean tracklength of 100, fill in 1e-2. Once the program is done, the table will be updated with the new tracking radii.

#### 2.6 Visualization of tracks

##### 2.6.1 Plotting options

**Show spots** Shows all detected spots in the current frame

###### Plotting tracks

- Show tracks in Range: Tracks are visible up to the displayed frame number. The parameter "#frames track is visible" defines how long tracks are plotted after the end of the track. A value of 0 will only show tracks visible in the current frame, for "inf" all tracks are plotted until the current frame.
- Show all Tracks: All tracks of the current movie are plotted
- Show initial positions: Only the position of first appearance of a track will be plotted

**Show events < shortest Track** All positions of non-linked detections are marked with yellow dots.

###### Coloring of tracks

- Random colored: The track color for each track is chosen randomly and the amount of colors can be entered in the field "#track-colors".
- Colored by track length: Matlabs "parula" color map is used to plot tracks with a color corresponding to their length. Shortest tracks are colored blue while longest tracks are colored in yellow.
- Colored by track length regime: The track colors are chosen according to user defined track length classes. Tracks with a minimum duration of the value entered in the field *min. length to count as long track* are plotted green whereas shorter tracks are shown in red.
- Colored by mean displacement: Matlabs "parula" color map is used to plot tracks with a color corresponding to their mean displacement. Tracks with the lowest mean displacement are colored blue while tracks with the highest mean displacement are colored in yellow.

##### 2.6.2 Plot selected tracks

Single tracks can be selected via the *Data Tips* function located in the toolbar of the main GUI. A click on "Plot selected track" in the main GUI opens a Matlab figure containing three plots (see Fig 2.6):

**Position plot** Plot of the positions of the whole track with track segments color coded by their intensities.

**Mean intensity plot** Shows the mean intensity inside a 3x3 pixel window around the spot center for each frame of the track. The plot is continued for 20 additional frames after the track ended (drawn in black) to check whether the fluorescent molecule has bleached or if the track was lost during tracking process.

**Kymographs** Opens a horizontal kymograph (YT, X-projection) and a vertical kymograph (XT, Y-projection) of the track. Here only one dimension is plotted at each timepoint while the other dimension is maximum projected within a window of 13 pixels around the spot center. Kymographs are continued for 20 additional frames after the track ended. A red line indicates the end of the track.

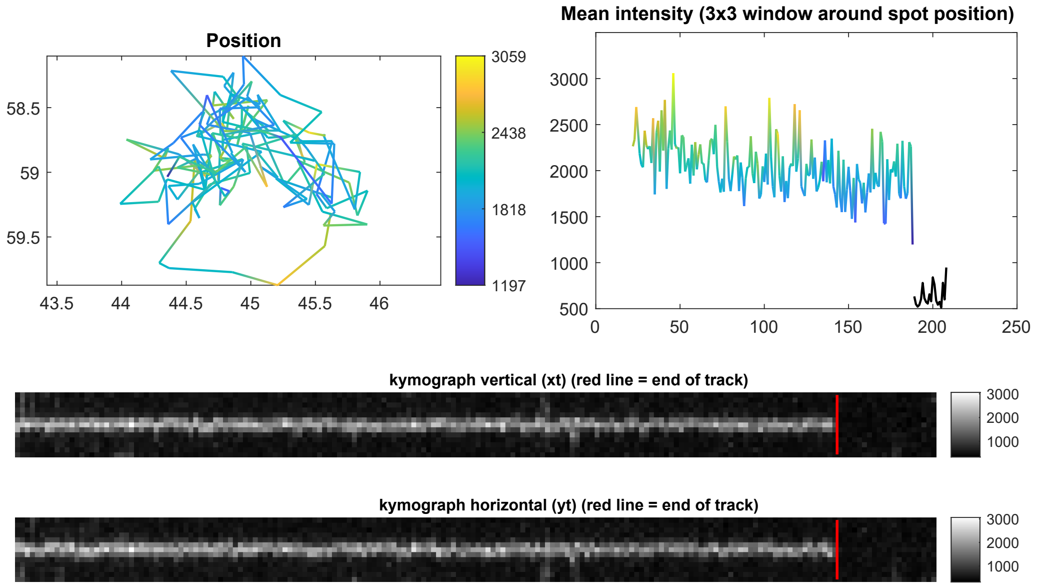

Figure 2.6: Track plotting tool.

##### 2.6.3 Detection mapping and jump distance mapping

Detected spots fit to sub-pixel precision with a 2D Gaussian fit are used to create a super-resolved heat map of localizations. Here, fitted positions of all spots are accumulated in a 2D histogram. The pixel values therefore correspond to the amount of detections in each pixel.

Similar to localization mapping, a heat map of jump distances can be created. For all jumps within a track, a virtual line is drawn between the start and end position of a jump. Each pixel touching this line is assigned with the corresponding jump distance. The resulting 2D histogram is then normalized by the amount of events in each pixel. For both kinds of heat maps the bin size can be chosen as a multiple of the original pixel size resulting in an up –or down-scaled image and can be entered in the field "Scaling factor".

#### 2.7 Track analysis tool

The track analysis tool can be accessed via "Tools" » "Track analysis tools".

##### 2.7.1 Overview

1. Batches analyzed and saved with the TrackIt software can be loaded into the track analysis tool via the button "Load batch .mat file(s)". The window below shows a list of loaded batch files where batches to be included in the analysis can be selected. Multiple batches can be selected using the "Ctrl" key.

2. The frame-cycle time selection window in the central left lists all frame-cycle times that are involved in the selected batches and can be used to select subsets of movies. Either a single movie, or a certain frame-cycle time or all movies can be selected to be included into the analysis.

3. The parameter selection window lists all the analysis parameters which are grouped into three tabs: mobility, tracked fractions and statistics. The selected parameter is then displayed in the central plot.

4. Shows a number of plotting options and settings regarding calculations of the different parameters:
   - #bins: Amount of bins in a histogram.
   - Jumps to consider: The maximum number of jumps that are considered in each track. This can be useful to prevent an over representation of slow diffusing or bound molecules in the histogram.
   - Normalization: Histogram normalization can be switched between "Count" and "Probability". Bin heights are either given by the number of events in each bin or normalized so that the height of all bins sums up to 1, respectively.
   - Units: Can be switched between "pixels & frames" and "microns & sec". If "pixels & frames" is selected, all results concerning length scales are given in the unit of pixels and all results concerning temporal units are in the unit of

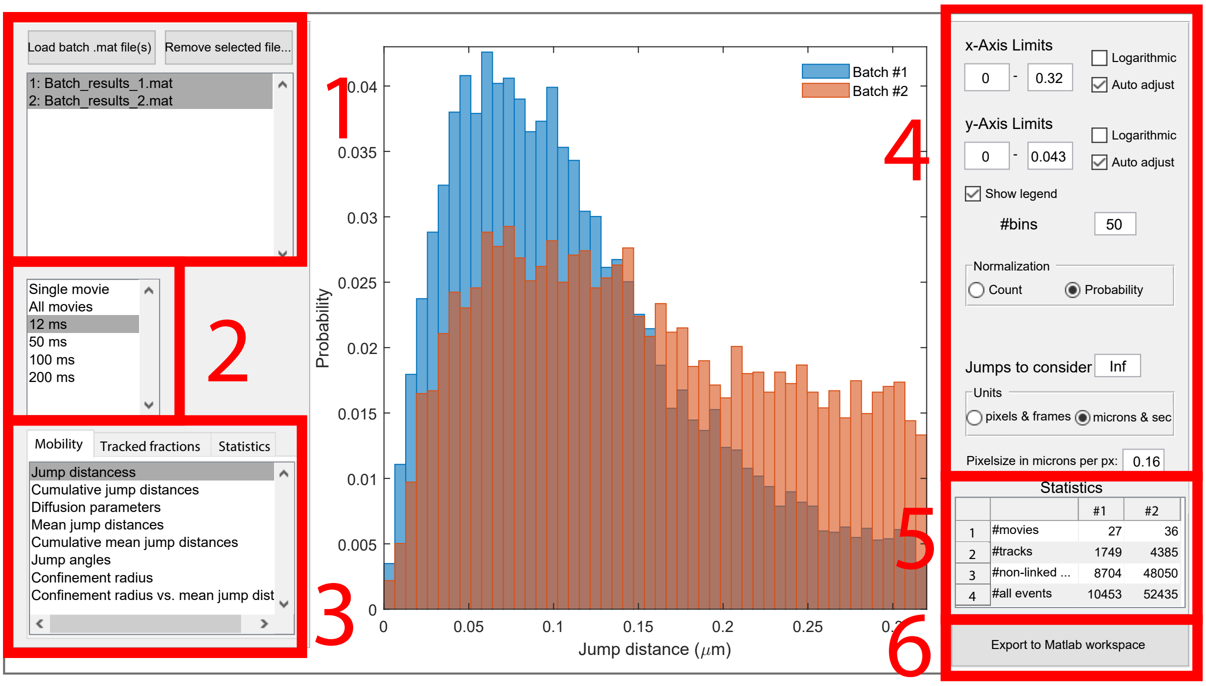

Figure 2.7: Screenshot of the track analysis tool. 1) Batch selection 2) Frame-cycle-time selection 3) Analysis parameter selection window 4) Plotting and analysis options 5) Statistics overview of loaded batches 6) Export to Matlab workspace.

frames. If "microns & sec" is selected, a conversion factor for the size of one pixel in $\mu m$ must be entered. All results are then displayed in micrometers using this conversion factor and seconds using the frame-cycle time stored for each movie (displayed in the frame-cycle time window).

- Normalize by ROI-size: Most of the statistical evaluations can be normalized by the size of the ROI in each movie in order to give information on densities (eg. spot densities).
- Illumination pattern: Can be switched between "periodic" and "ITM" (interlaced time-lapse microscopy). ITM is an illumination scheme where two subsequent image acquisitions are followed by a dark time. This scheme is specifically designed to gather quantitative information on chromatin-bound fractions and proportions of stable bound molecules [11]. Due to a non-periodic image acquisition, the illumination scheme has to be taken into account when calculating the number of dark periods that an immobile molecule survived (see Section 2.7.3).

5. The statistics overview windows gives information on movie numbers and molecule counts for each loaded batch.

6. Exports two variables to the Matlab workspace: "currentPlotValues" contains the

data which is shown in the current plot, "allHistogramResults" contains all the raw data that is used to display the different parameters implemented in the histogram tool.

##### 2.7.2 Mobility analysis tab

**Jump distances** Displays the distribution of jump distances in a histogram. The jump distance is defined as the Euclidean distance between the positions of two linked spots of a track in consecutive frames.

**Cumulative jump distances** Displays the cumulative distribution of squared jump distances. This distribution can be used to fit either a 2-rate or 3-rate diffusion model (see below). The fitted diffusion functions can be visualized by selecting "show 2-rate diffusion fit" and "show 3-rate diffusion fit".

**Diffusion parameters** Cumulative distributions of squared jump distances are fitted with two or three exponential components corresponding to two or three effective diffusion constants $D_{1-2}$ or $D_{1-3}$ with amplitudes $A_{1-2}$ or $A_{1-3}$, respectively [1]. If the two-rate model is selected in the pop-up menu, the following fit function is applied

$$f_2(X) = A_1\left(1 - e^{-\frac{X}{D_1}}\right) + (1 - A_1)\left(e^{-\frac{C_1}{D_2}} - e^{-\frac{X}{D_2}}\right) / \left(e^{-\frac{C_1}{D_2}} - e^{-\frac{C_2}{D_2}}\right) \tag{2.1}$$

If the three-rate model is selected in the pop-up menu, the applied fit function is

$$\begin{aligned} f_3(X) = & A_1\left(1 - e^{-\frac{X}{D_1}}\right) + A_2\left(1 - e^{-\frac{X}{D_2}}\right) \\ & + (1 - A_1 - A_2)\left(e^{-\frac{C_1}{D_3}} - e^{-\frac{X}{D_3}}\right) / \left(e^{-\frac{C_1}{D_3}} - e^{-\frac{C_2}{D_3}}\right) \end{aligned} \tag{2.2}$$

Functions are normalized to account for the lower and upper limit of jump distances $C_1 = 0$ and $C_2 = d_{max}$ where $d_{max}$ is the tracking radius. Error bars in the plot indicate 95% confidence intervals of the fitted diffusion constants and amplitudes. To ensure that the fit converges to a global minimum, diffusion constant start values can be entered on the right side of the figure. For a visualization and control of the fitted function see *cumulative jump distances* above.

Important: fitting distributions populated with jump distances from mixed frame-cycle times will lead to wrong results!

**Mean jump distances** Shows a histogram of the mean jump distances of all tracks. The mean jump distance $\bar{d}$ of a track is defined as:

$$\bar{d} = \frac{1}{n}\sum_{i=1}^{n} d_i \tag{2.3}$$

where n is the number of jumps in a track and $d_i$ are the jump distances of the track.

**Cumulative mean jump distances** Displays a cumulative histogram of the mean jump distances.

**Jump angles** Displays a polar histogram of jump angles which are defined as the change in direction between each step of a track. The jump angle histogram can give additional information about diffusion properties or directional behavior. A circular polar histogram may indicate a random, brownian motion. A polar histogram where bins are concentrated around 180° indicates either a confined diffusion or a bound state (where the apparent displacement due to a localization imprecision is higher than the fluorescent molecule movement). Tracks with its polar histogram centered near 0° tend to have only small directional changes and may indicate a directed motion. See also [6] and [2].

**Alpha values from MSD fit** Shows the distribution of alpha-values which are extracted by fitting the mean squared displacement (MSD) with a power law

$$MSD = 4 \cdot D \cdot t^{\alpha} \tag{2.4}$$

where $D$ is the diffusion coefficient and $\alpha$ a coefficient that indicates the motion type. The MSD is calculated as

$$MSD(\tau) = \frac{1}{\tau}\sum_{t=1}^{\tau}(x(t) - x(t+\tau))^2 + (y(t) - y(t+\tau))^2 \tag{2.5}$$

where x(t) and y(t) are the coordinates of the spot within a track at the time t. $\tau$ is the maximum time interval between two spots of a track and can be set in the field "MSD cut-off". A minimum track length can be entered in the field "shortest track".

**Diffusion constants from MSD fit** Shows the distribution of diffusion constants which are extracted by fitting the MSD (see above).

**Confinement radius** Shows the distribution of confinement radii calculated by fitting the MSD with a confined diffusion model [8]

$$MSD_{circle} = R^2 \cdot \left(1 - e^{-4 \cdot D \frac{t}{R^2}}\right) + \text{offset} \tag{2.6}$$

where R is the radius of confinement and D is the local diffusion coefficient. The offset is introduced to account for the finite localization precision. To use exclusively tracks which show a confined motion, tracks with alpha-values above a threshold specified in the field "upper alpha limit" are discarded.

**Confinement radius vs. mean jump distance** Displays a 2-dimensional plot where the mean jump distance of a track is plotted versus its confinement radius (see above) [8]. This representation can give insights into different mobility classes of single molecules.

##### 2.7.3 Tracked fraction analysis tab

**Bound fractions (Bf)** Bound fractions can either be analyzed from movies with continuous or interlaced illumination. The illumination pattern can be selected in the "Illumination pattern" button group. The fractions of molecules belonging to two different binding time classes can be approximated using the interlaced time-lapse microscopy (ITM) scheme [4, 11]. In ITM, two frame acquisitions separated by a short dark time are followed by a longer dark time. This illumination scheme allows to classify molecules by their binding time. Molecules count as long bound ($N_{\text{long}}$) if they survive at least one long dark time, short bound ($N_{\text{short}}$) if the molecule survives only one short dark time and diffusive if the molecule is detected in only one frame ($N_{\text{non-linked}}$). The ITM scheme can be used for a bound fraction analysis by choosing the option "Interlaced (ITM)" in the field "Illumination pattern". Choosing "Continuous", bound fractions can also be calculated from continuous movies, but come with high errors and have to be handled with care. Generally, accurate bound fractions can only be obtained if bleaching is corrected [11]. Three different types of bound fractions can be extracted:

**Tracks vs. all events** Shows the fraction of all bound molecules with respect to all counted events

$$\mathrm{Bf}_{\text{all bound}} = \frac{N_{\text{bound}}}{N_{\text{all events}}} = \frac{N_{\text{long}} + N_{\text{short}}}{N_{\text{long}} + N_{\text{short}} + N_{\text{non-linked}}} \tag{2.7}$$

where $N_{\text{bound}}$ is the number of all molecules classified as bound and $N_{\text{all events}}$ is the number of all events.

**Long tracks vs. all events** Shows the fraction of long bound molecules with respect to all counted events

$$\mathrm{Bf}_{\text{long vs. all}} = \frac{N_{\text{long}}}{N_{\text{all events}}} = \frac{N_{\text{long}}}{N_{\text{long}} + N_{\text{short}} + N_{\text{non-linked}}} \tag{2.8}$$

where $N_{\text{all events}}$ is the number of all events.

**Long tracks vs. long + short tracks** Shows the fraction of long bound molecules with respect to all bound molecules

$$\mathrm{Bf}_{\text{long bound}} = \frac{N_{\text{long}}}{N_{\text{bound}}} = \frac{N_{\text{long}}}{N_{\text{long}} + N_{\text{short}}} \tag{2.9}$$

**Pooled and movie-wise values** Bound fractions are determined for each movie (displayed as blue triangles) and can show a significant variance between cells or movies. The average of these movie-wise values is calculated and displayed as a red dot. To avoid an over-representation of outliers or movies with low molecule counts, events for each binding time class can be summed up over all movies resulting in a single "pooled" bound fraction of all movies which is displayed as a black dot.

**Error estimation** The error for the movie-wise bound fraction (red error bar) is given by the standard error of the mean

$$\Delta_{\mathrm{BF}_{\text{moviewise}}} = \frac{\mathrm{stdev}(\mathrm{BF}_{\text{moviewise}})}{\sqrt{N_{\text{movies}}}} \tag{2.10}$$

where $\mathrm{stdev}(\mathrm{BF}_{\text{moviewise}})$ is the standard deviation of the movie-wise bound fraction values and $N_{\text{movies}}$ is the amount of movies for which bound fractions are calculated.

The error for the pooled bound fraction (black error bar) is calculated through linear error propagation

$$\Delta_{\mathrm{BF}_{\mathrm{pooled}}} = \frac{1}{N_{\mathrm{denominator}}} \cdot \delta_{N_{\mathrm{numerator}}} + \frac{N_{\mathrm{numerator}}}{N^2_{\mathrm{denominator}}} \cdot \delta_{N_{\mathrm{denominator}}} \tag{2.11}$$

where $N_{\mathrm{numerator}}$ is either $N_{\mathrm{long}}$ or $N_{\mathrm{bound}}$ and $N_{\mathrm{denominator}}$ is either $N_{\mathrm{bound}}$ or $N_{\mathrm{all\ events}}$. The errors $\delta$ are estimated by the counting errors with $\delta_{N_{\mathrm{numerator}}} = \sqrt{N_{\mathrm{numerator}}}$ and $\delta_{N_{\mathrm{denominator}}} = \sqrt{N_{\mathrm{denominator}}}$.

**No. of tracks** Shows the total number of tracked molecules in each movie.

**No. of non-linked detections** Shows the number of non-linked detections. This can either be single detections that have not been linked to tracks or detections of a track with a track length shorter than the minimum track length set by the user (see Fig. 2.1 panel 4).

**No. of all events (tracks + non-linked)** Shows the number of total events in each movie calculated by the total number of tracks plus the number of non-linked detections.

**No. of long tracks** In the case of continuous illumination the amount of tracks which are longer than the threshold entered in the field "count as long track if track is longer than" is shown. If the interlaced illumination pattern (ITM) is selected, the displayed values are given by the amount of tracks which survive a specific number of long dark periods entered in the field "#survived dark periods to count as long track".

**No. of short tracks** Shows the amount of short tracks in each movie which is given by the total number of tracks minus the number of long tracks (see above).

##### 2.7.4 Statistics tab

**Track lengths** Histogram showing the lengths of tracks. The track length is defined as the amount of frames a track survives. The minimum length of a track is therefore two.

**Avg. track length** Shows the average track length in each movie.

**Avg. no. tracks per frame** The average number of tracks per frame in a movie is calculated by counting the number of tracks visible in each frame divided by the number of frames in the corresponding movie.

**Avg. no. spots per frame** Shows the average number of spots per frame in each movie calculated by dividing the total amount of spots by the number of frames in the movie. If "Normalize by ROI-size" is selected, the values are divided by the size of the ROI resulting in the average spot density per frame. Based on experience, the average spot density should be below $2.5 \cdot 10^{-3} \frac{\text{spots}}{\text{px} \cdot \text{frame}}$ for the nearest neighbor algorithm to link trajectories correctly.

**ROI size** Shows the size of the region of interest (ROI) of each movie either in $\mu m^2$ or in number of pixels depending on the selected units.

#### 2.8 Analysis of dissociation rates with GRID

Via "Tools" » "Analyse dissociation rates (GRID)" tracking results can be directly analyzed using the GRID software. Therefor track lengths (in seconds) are passed over to the GRID tool. For a detailed description of GRID please read the GRID user manual or refer to [10]. GRID is fully compatible with batch files created with the TrackIt software and can be loaded directly from the GRID GUI via "Files" » "Load File". GRID is available under `https://gitlab.com/GebhardtLab/GRID`.

#### 2.9 Data export and movie creation

##### Export all data to Matlab workspace

By selecting "File" » "Export all data to Matlab workspace" in the menu bar of the main GUI, the data structure is saved in a variable of type "struct" with the name "currentBatch" in the Matlab workspace.

##### Export tracks

An export dialog window (see Fig 2.8) can be opened by clicking "File" » "Export tracks". In the "Compatibility" popup-menu .mat files can be chosen to be compatible either with Spot-On [3] or with vbSPT [9]. Only .csv files are compatible with Spot-On. Spot-On is available under `https://spoton.berkeley.edu/` and vbSPT under `https://github.com/bmelinden/vbSPT`.

##### Movie creation

A movie inlcuding all selected plot elements can be created by clicking "Tools" » "Create .avi from current movie". A .avi movie file will be created using the exact same plot

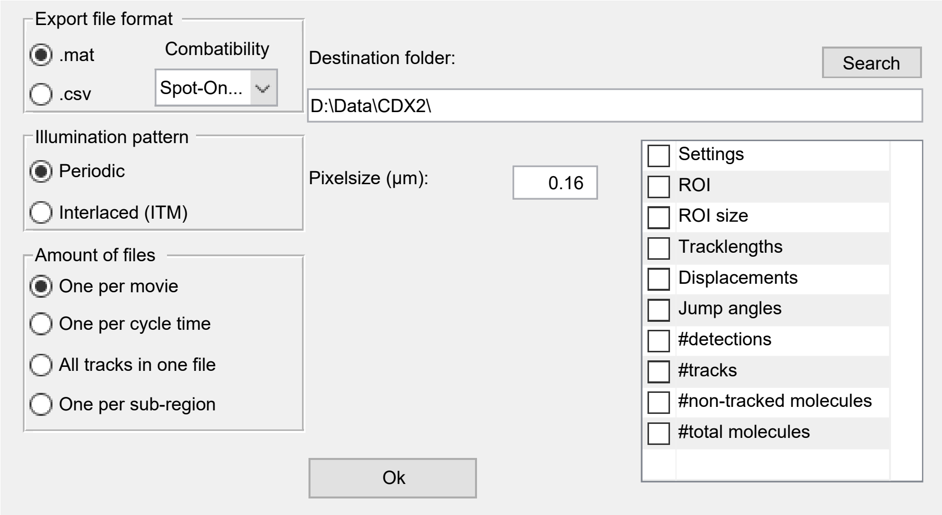

Figure 2.8: Track export window.

properties as currently shown in the main GUI. The playback speed (frames-per-second, FPS) can be set in the "FPS" field at the lower part of the main GUI.

#### 2.10 Additional tools

##### 2.10.1 Histogram of spots

A tool showing information about the detected spots can be open via "Tools" » "Histogram of spots" where the following data can be shown and retrieved:

**Peak spot intensity** Shows a histogram of the peak intensity of all detected spots. The peak intensity is defined as the highest pixel value of all pixels within a radius of 2 px around the spot center.

**Fitted spot intensity** Shows a histogram of the fitted intensities of all detected spots as given by the maximum of the 2D Gaussian fit.

**Spot SNR** Histogram of the signal-to-noise ratio (SNR) of all detected spots. The SNR is calculated in a square window with side length of 17 pixels around the spot peak position at the center pixel (see Fig 2.10) as follows:

$$\mathrm{SNR} = \frac{\overline{I}_{\mathrm{blue}} - \overline{I}_{\mathrm{white}}}{\sigma_{\mathrm{white}}} \tag{2.12}$$

where $\overline{I}_{\mathrm{blue}}$ and $\overline{I}_{\mathrm{white}}$ is the mean intensity of the pixel values of the respective area and $\sigma_{\mathrm{white}}$ denotes the standard deviation of the the pixel values of the white area.

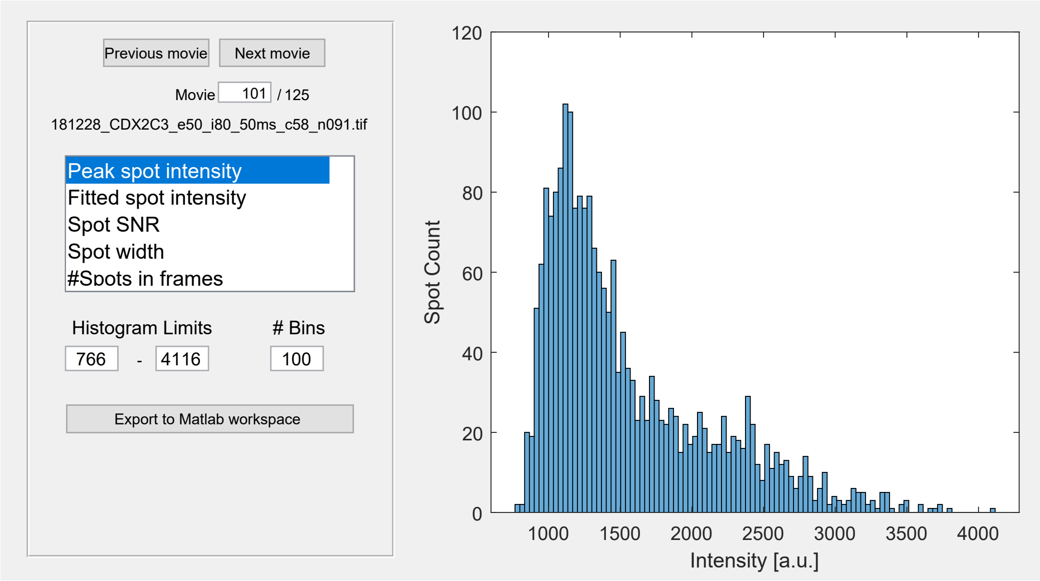

Figure 2.9: Spot histogram tool.

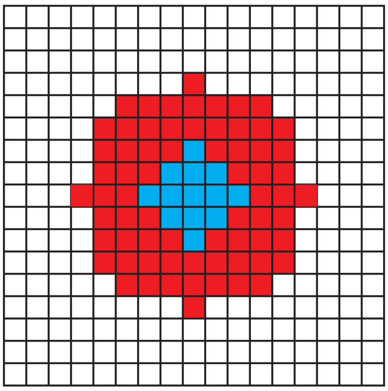

Figure 2.10: Pixel mask used for calculating the spot signal-to-noise ratio (SNR). **White**: Area where the mean background signal $\overline{I}_{\text{white}}$ and standard deviation of background noise $\sigma_{\text{white}}$ are calculated. **Blue**: Area where the mean intensity $\overline{I}_{\text{blue}}$ of the spot is calculated. **Red**: Not used for calculation

**Spot width** Histogram of the spot width of all detected spots as given by the standard deviation $\sigma$ of the Gaussian fit.

**#spots in frames** Shows a plot of the number of detected spots in each frame.

##### 2.10.2 Movie splitter

Movies containing dark frames or movies containing more than one channel, as commonly originated from multiple-color experiments, can be split using the "Movie splitter" tool located in the "Tools" menu bar (see Fig 2.11). The amount of splits as well as the number of frames in each sequence can be defined by the user. A name entered in the field "Add-on to original filename" will be added to the original filename for each splitted movie part. If desired a .txt file containing the .tiff metadata can be created for each original movie. TrackIt uses the Bio-Formats library (`https://www.openmicroscopy.org/bio-formats/`) to extract metadata.

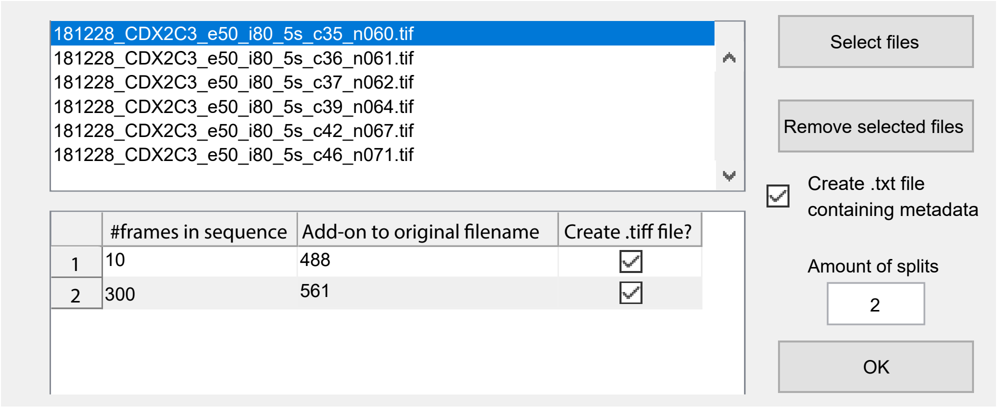

Figure 2.11: Movie splitter tool.
